# A Geophysical and Climatological Assessment of New Guinea — Implications for the Origins of *Saccharum*

**DOI:** 10.1101/2020.06.20.162842

**Authors:** Dyfed Lloyd Evans

## Abstract

Any assessment of whether or not *Saccharum* species are native or introduced in New Guinea require an evolutionary (in a geological sense), geophysical and climatological assessment of the island. Like many of the land masses circling the Pacific (in the volcanically active region known as the ‘ring of fire’) New Guinea is geologically young, with the island in its modern form not pre-dating 2 Ma. Novel modelling of the 74 ka youngest Toba supereruption indicates a potential extinction level tsunami and loss of habitat. The late Pleistocene megafaunal mass extinction and the last glacial maximum (33–16 ka) are two global effects that would have significantly altered the flora on New Guinea; though the implications of these events on New Guinea have not previously been studied. Even if the genus *Saccharum* was established on the island during pre-historic times the consequences of Toba and other global climate change events means that it would have been eliminated from New Guinea and would have had to be re-introduced during the period of human colonization. Indeed, given the evolution of *Saccharum*’s immediate ancestors in Africa and Indochina it is most parsimonious to conclude that it was never native to New Guinea, but was introduced by humans relatively recently.

Little work has been done on palaeotsunami evidence and ancient tsunami modelling in New Guinea. However, the recent recognition that the Aitape skull (dating to about 6 ka) may have been the victim of a tsunami (Goff et al. 2017) show that, in the past, tsunami have pen etrated significantly (about 10 km in this case) into the interior of the island to have a profound effect on biodiversity. This tsunami would have left the north coast of the island impoverished of plant life for several decades after.

## Introduction

The early evolutionary history of sugarcane is intimately linked to the geophysics of the region(s) where sugarcane is native. Extending back from modern *Saccharum* hotspots this area of interest must extend from Indochina through Wallacea and into Melanesia.

At the time where models of sugarcane origins were being developed (principally the 1940s and 1950s) little was known or accepted about plate tectonics (the theory was not generally accepted until 1956 (Runcorn 1956; Carey 1956)). The site of lake Toba in Sumatra as the locus of a major supervolcano eruption had not been recognized (Zelinski et al. 1996) and little was known about glaciation and its effects (see later in the tex). Modern theories on linguistic evolution had not been developed (see main text) and theories on Melanesian and Austranesian colonization were in their infancy. The effects of humans on transporting flora and fauna outside their native ranges and the use of agriculture to create centres of diversity had not yet been fully recognized.

In terms of the origin of sugarcane there are two competing theories. The first is that sugarcane evolved naturally on the Indochinese mainland and/or arose by natural hybridization of *S*. *officinarum* and *S*. *spontaneum*. The second, and most pervasive, hypothesis is that sugarcane arose by human selection from a purported ancestral species, *S*. *robustum* in New Guinea. To properly compare the evidence for the two hypotheses, we need to assess the viability of New Guinea as a locus for the evolution of *Saccharum* and sugarcane.

It should be noted also that whilst the evidence for an early inception of agriculture in New Guinea is unequivocal — with an even earlier phase of plant mmanagement in the natural environment — it should be noted that the evidence for the presence of sugarcane as an early crop is vanishingly small (Denham 2011).

### On the Origins of New Guinea

Like many of the island chains circling the pacific plate faults, the geology and formation of New Guinea is relatively recent. New Guinea lies on the northern boundary of the Australian plate and sits on a stable platform of Paleozoic crystalline rocks about 150 km wide (Dow 1977). This is overlaid by Mesozoic and Paleogene sedimentary rock. During the Paleogene, a major up-faulting and resultant island arc volcanism occurred. Volcanism subsided until the late Eocene/early Oligocene when increased plate movement created low-grade metamorphic rock along the length of the mobile belt. During the Miocene limestone was deposited on the ocean shelfs. Though the main, central, part of New Guinea is an ancient part of the Australian plate, the whole mass of New Guinea probably arose by multiple island collisions across the main fault line. These occurred in multiple docking stages from 25 Ma to 2 Ma with the final stages forming the characteristic Bird’s Head and Bird’s Neck part of the island. It is these collisions that gave rise to the basins, high mountains, and isolated valleys that are the characteristic features of New Guinea (Pigram and Davies 1987). It is not conceivable that any ancestral *Saccharum* species or species ancestral to *Saccharum* could have colonized New Guinea, and survived on the island, prior to this date.

In geological time, New Guinea is a relatively recent land mass, having matured within the past 2 million years. This also seems to agree with the time at which the island was first colonized by many species (Toussaint et al. 2014; Oliver et al. 2017).

### Potential Extinction Events within the Timescale of Human Settlement

New Guinea represents one of the sites in the world where anatomically modern humans arrived and settled earliest. Estimates for settlement range from 60–50 ka (O’Connell & Allen 2004; Summerhayes et al. 2010). In these terms, the major extinction events that lie in this timescale are the Toba supervolcano eruption (73.88±0.32 ka) (Storey et al. 2012) which just pre-dates human settlement, the late Pleistocene megafaunal extinction (c. 45 ka) and the last glacial maximum (33–16 ka). As these events would have had major effects on the flora of New Guinea, they are each discussed/modelled in this paper.

### New Guinea and a Tsunami-induced Inundation

As the major tuff dispersal from the Toba supervolcano eruption was largely in a westerly/south westerly direction (Figure 1), little work has been performed to analyze potential Toba-induced tsunami waves outside the Indian ocean. In truth, with many straits and narrow waterways in the vicinity of Sumatra, particularly the Malacca strait, these could allow for megatsunami propagation towards New Guinea. As such a potential large-scale event in this direction has not been previously modelled it is examined in detail in this paper.

**Figure 1:**
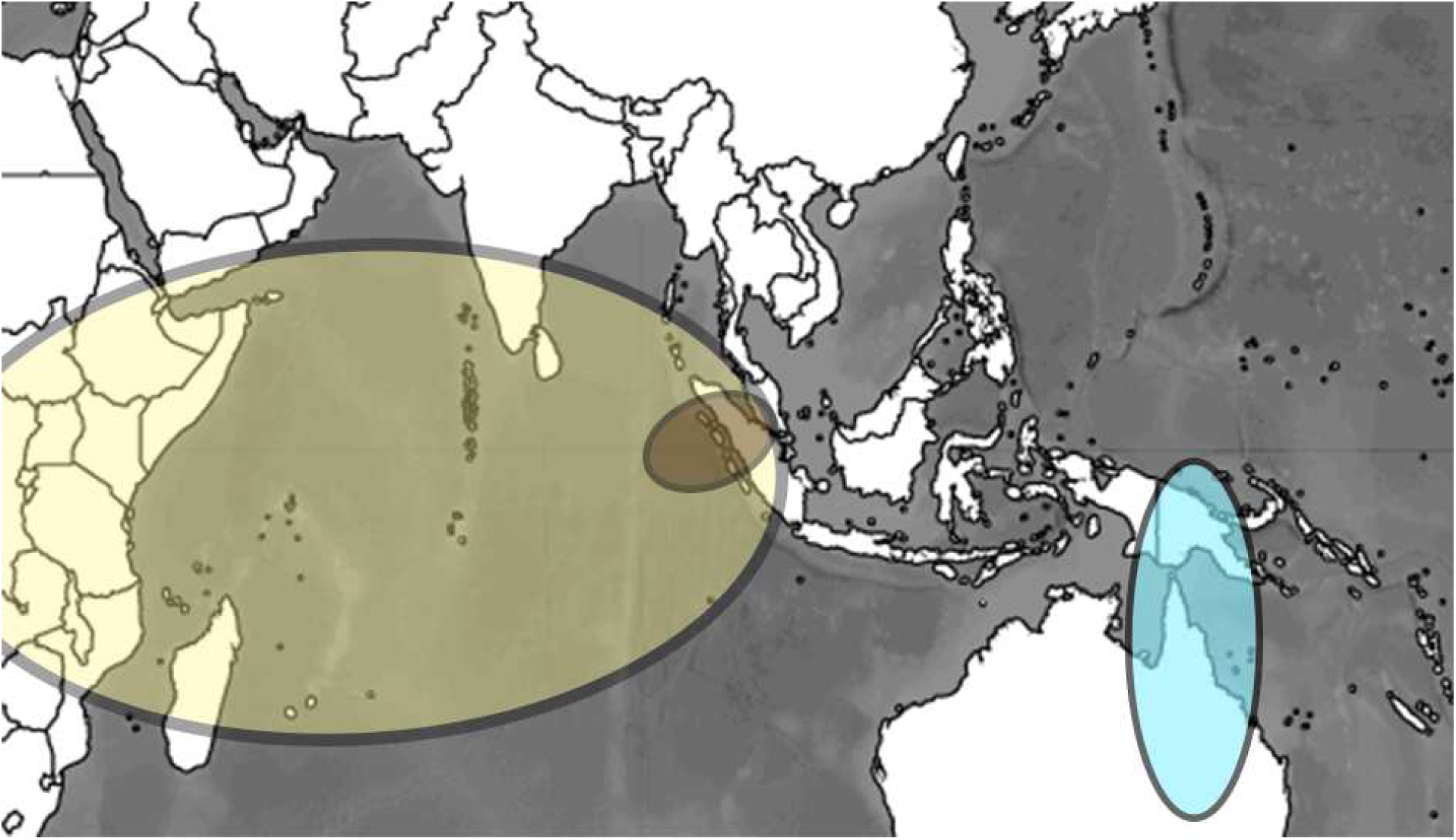
Placing the Toba eruption on a map of the Western Pacific. The extent of the youngest Toba tuff deposits are shown in yellow. In brown is the extent of the 100m deposit from Toba. Also shown is a tuff deposit from an eruption off the coast of New Guinea occurring 45–90 ka (Coulter et al. 2009) (shown in blue).

Tsunami essentially have three phases: inception, propagation and inundation. The inception phase is the trigger for the tsunami, be that undersea earthquakes, landslides or volcanic eruptions. Propagation is the way the wave travels through the water and the mechanisms by which oceanic depth and beach topology affect the energy and height of the wave. The final phase is when the wave makes landfall. Mathematically, tsunami waves have been extensively studied and the fluid dynamic equations describing tsunami have also been applied to pyroclastic flows and tuff deposition from volcanic eruptions (Costa et al. 2014). Unlike modelling historical tsunami events, in the case of the Toba supervolcano eruption eruptive extents have had to be modelled backwards from existing evidence.

This becomes all the more necessary as recent modelling of Toba dense rock equivalent (DRE) ejecta (Costa et al. 2014) indicates a 100m depth of DRE adjacent to the main eruption site. Though the main direction of the major DRE fall was south-west of Sumatra into the Indian Ocean, the 100m DRE also impacted the strait of Malacca. Core samples measured by Matthews et al. (2012) demonstrated a tephra depth of 150 cm in Tampan, Malaysia (382 km from the Toba vent). This is good evidence that the Malacca strait would have been inundated with ejecta from the Toba event. Under these circumstances tsunami are almost inevitable. With DRE falling into the Malacca strait the water would have been displaced and would have flowed northwards into the Andaman sea and southwards into the Java sea. It is this southerly flow and displacement of oceanic water that concerns this analysis.

### Biogeography of New Guinea

Biogeographically, New Guinea is part of Australasia rather than the Indomalayan realm and the fauna is overwhelmingly Australian. However, botanically, New Guinea is considered part of Malesia, a floristic region that extends from the Malay Peninsula across Indonesia to New Guinea and the East Melanesian Islands. The flora of New Guinea is a mixture of many tropical rainforest species with origins in Asia, together with typically Australasian flora. Results suggest that species-level diversification within New Guinea has been recent (*<*5 Ma), corroborating geological evidence that dates substantial landmass formation to *<*10 Ma (Hall 1998; Cloos 2005) with final stabilization of the island at ∼2 Ma (Pigram and Davies 1987). Current models suggest that that New Guinea was largely submerged until the Early Pliocene and that formation of the present large emergent area occurred in the past 5 Ma, with collisions of smaller islands between 5 Ma and 2 Ma forming the final land mass, as we know it today.

Work on invertebrate evolution in New Guinea (Touissant et al. 2014) suggests recent colonization of the island about 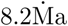 possibly with an origin in the central orogen and early diversification at montane level with several altitudinal shifts. The authors argue that ‘Strong landscape changes during the uplift as well as significant amounts of volcanism during the late Neogene followed by Quaternary climate change likely shifted environmental conditions.’ They also found evidence for only two very recent colonizations into the Bird’s Head region (which was the final part of New Guinea to evolve, palaogeographically about 2 Ma.

### On the Origins of Genus Saccharum

The geological age of New Guinea at its first habitable stage is just congruent with the time of origin of extant *Saccharum* species. Indeed, our data (Lloyd Evans & Joshi 2016; Lloyd Evans et al. 2019; Lloyd Evans 2020) indicate that *S*. *spontaneum* (wild sugarcane) evolved from a common ancestor with *Narenga porphyrocoma about 1*.*5 million years ago*. *The* Narenga*+*Saccharum *clade shared a common ancestor with African* Miscanthidium species about 2.2 Ma. *Miscanthidium* is today found in central and eastern Africa down to south Africa. Whilst *Narenga* is found in India and Indonesia with a rump population in Ethiopia. Comparison of chloroplast and genomic data indicates that *Narenga* and *Saccharum* hybridized after the two genera split from each other, indicating co-localization after the divergence event 1.5 Ma (Lloyd Evans, 2020). Narenga is not found in New Guinea, nor is the more ancestral *Miscanthidium* genus. For their initial inception, the origins of *Saccharum* species lie in the spread of the African C_4_ savanna.

This firmly places the origins of the Saccharinae (the subtribe that includes *Saccharm* and all its close relatives) in Africa, and their subsequent evolution into genus *Saccharum* in Indochina. Indeed, of the seven genera that comprise the Saccharinae (Lloyd Evans and Hughes 2020) half are native to Africa. By 0.64 Ma all major *Saccharum* species had evolved (Lloyd Evans and Joshi, 2016). However, if they had reached Malesia the local effects of the Toba eruption would almost certainly have eliminated them from the region.

## Results and Discussion

### Toba Tsunami Modelling — Initial Conditions

The work of Costa et al. (2014) indicated a 100m depth of dense rock equivalent (DRE) material landing in the strait of Malacca. Their model indicates a 250km length for the extent of rockfall into the strait. The strait is an average of 60km wide at the site where the Toba ejecta fell. Combining all the information gives a DRE fall of 25000 × 60000 × 100 = 1.5 × 10^12^*m*^3^ of DRE material falling into the Malacca strait. Employing a conservative estimate of DRE mass as 800kg/m^3^ (Wallace et al. 2010) yields a total mass of 1.2 × 10^15^kg. The kinetic energy of a falling body is 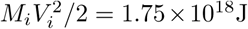 (at terminal velocity). Today, the south of the strait is shallow, rarely exceeding 25m, though it can be 100m in the shipping lanes (Amiruddin et al. 2011) but at the north of the strait depths can exceed 150m. Assuming that pre-Toba depths in the impact area were 250m then this gives the maximal amplitude for wave propagation (Ward & Asphaug 2003) and the initial conditions are derived.

Beyond wave initiation (inception) and propagation, handled in equations 1-5 of the Materials and Methods section, this leaves the final aspect of tsunami waves, inundation. Inundation represents the processes occurring when tsunami reach the edge of the land mass and traverse from the marine to the terrestrial environment. The major factors affecting the damage done by inundation are water depth in the run-up to the shore and the slope of the shore itself. The Banda Sea, which immediately precedes New Guinea has very deep waters (up to 7.2km). One of the issues affecting the propagation of tsunami waves onto shore is that Equation 5 is that *S*_*L*_(*ω*, **r**, **r**_*i*_) favours low-frequency waves. Typically this is reduced to Green’s law [*h*(**r**_1_)*/h*(**r**)]^1*/*4^ but this tends to ∞ as depth approaches zero. In shallow seas, and on shallowly shelving beaches Equation 5 tends to over-predict tsunami height near shore. For the case of the Toba-associated tsunami and New Guinea this is not necessarily the case as the Banda Sea is a deep ocean trench and New Guinea’s volcanic/fault line origins generally gives it a steeply-sloping shoreline, thus the usual correction is omitted here. For completeness, however, the run-up scaling factor is defined (Equation 10).

### Wave Height and Propagation

One of the major unknowns affecting wave heights as they approach the New Guinea coast is the depth of the Malacca straits. In the initial simulations, a 250m depth was chosen (modern sea floor + 100m depth of Toba ejecta). However, as the 2004 tsunami significantly affected water depth in the straits (Singh 2010) it is possible that ocean floor depths may have been closer to 1000m prior to the Toba supereruption. Thus modelling was performed on successive water depths in 250m increments up to 1km.

As the area in question is a strait and not open ocean the assumption of even dispersal of the cavity wave (Equation 3) is not sensible. Initially, all the energy of the impact would have been directed at the propagating wave front. Even if 60% of the available energy from impact caused water to splash over the sides of the strait and onto land, 20% additional total energy (the waves would have travelled in both directions along the channel) could increase initial wave heights by a factor of 2–3. This is assuming that only about 8% of the total energy of impact was converted into wave energy (this is conservative, as asteroid impacts are calculated to transfer almost 10% of total energy into wave energy (Ward & Asphaug 2003)).

Initial propagation of the wave front is shown in Figure 2, with intervals at 10 minutes. Based on a range of starting conditions, minimum and maximum wave heights are presented in Table 1.

**Table 1:** Table showing the effect of initial wave propagation heights on the maximal tsunami wave height making landfall at New Guinea. Γ is an empirical correction factor, taking into account that the initial wave was initiated in a strait and not the open ocean.

|  | Wave Height |  |  |  |
| --- | --- | --- | --- | --- |
|  | initial (above sea level)/landfall (max height) (m) |  |  |  |
| Depth of Malacca Straits (m) | $\Gamma = 1$ | $\Gamma = 1.2$ | $\Gamma = 1.4$ | $\Gamma = 1.6$ |
| 250 | 100/460 | 120/550 | 140/640 | 160/730 |
| 500 | 150/490 | 180/560 | 210/685 | 245/785 |
| 750 | 180/520 | 220/625 | 260/730 | 305/830 |
| 1000 | 220/580 | 265/700 | 310/810 | 345/930 |

**Figure 2:**
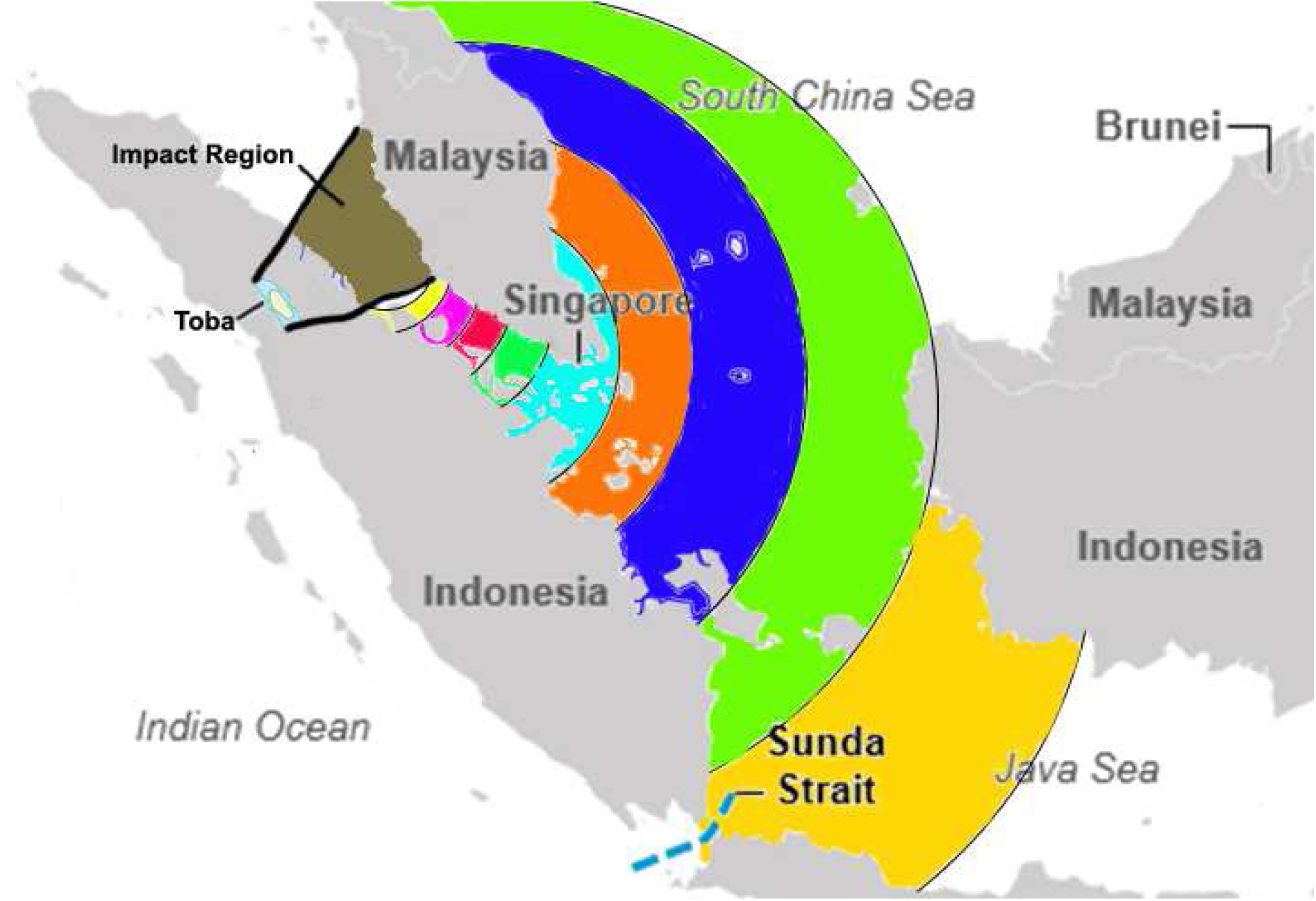
Initial propagation of the tuff fall tsunami wave from the Malacca straits. Colour bands show the propagation of the wave at 10 minute intervals from initial progression to entering the Java sea.

The left-hand column in Table 1 would be equivalent to a deep ocean asteroid impact. Increasing the depth of the Malacca strait increases the depth of water at which the initial wave starts, and increasing Γ also increases this starting wave height. Given the age of the Toba supereruption (74 ka) a depth between 750 m–1 km is likely. The closest real and modelled event to that analyzed in this paper is the 1958 Lituya Bay tsunami in Alaska. Here, a combined earthquake and landslide induced a 520 m high tsunami in an inlet with a maximal depth of 220 m and an average depth of 120 m. Modelling (Mader & Gittings 2002) indicated that maximal wave height above sea level was 250 m, with a run-up of 524m.

In many ways the modelling of the Toba tsunami impact on New Guinea reflects what was actually observed in the Lituya Bay tsunami. However, modelling the increase in amplitude of the initial wave gives increasingly large land-falls in New Guinea with runup wave heights of 600m to 1km predicted in the most likely median prediction area.

However, it’s not just wave heights that are important, it is the total flow of water and mapping the wave currents onto a topological map of New Guinea yields Figure 3.

**Figure 3:**
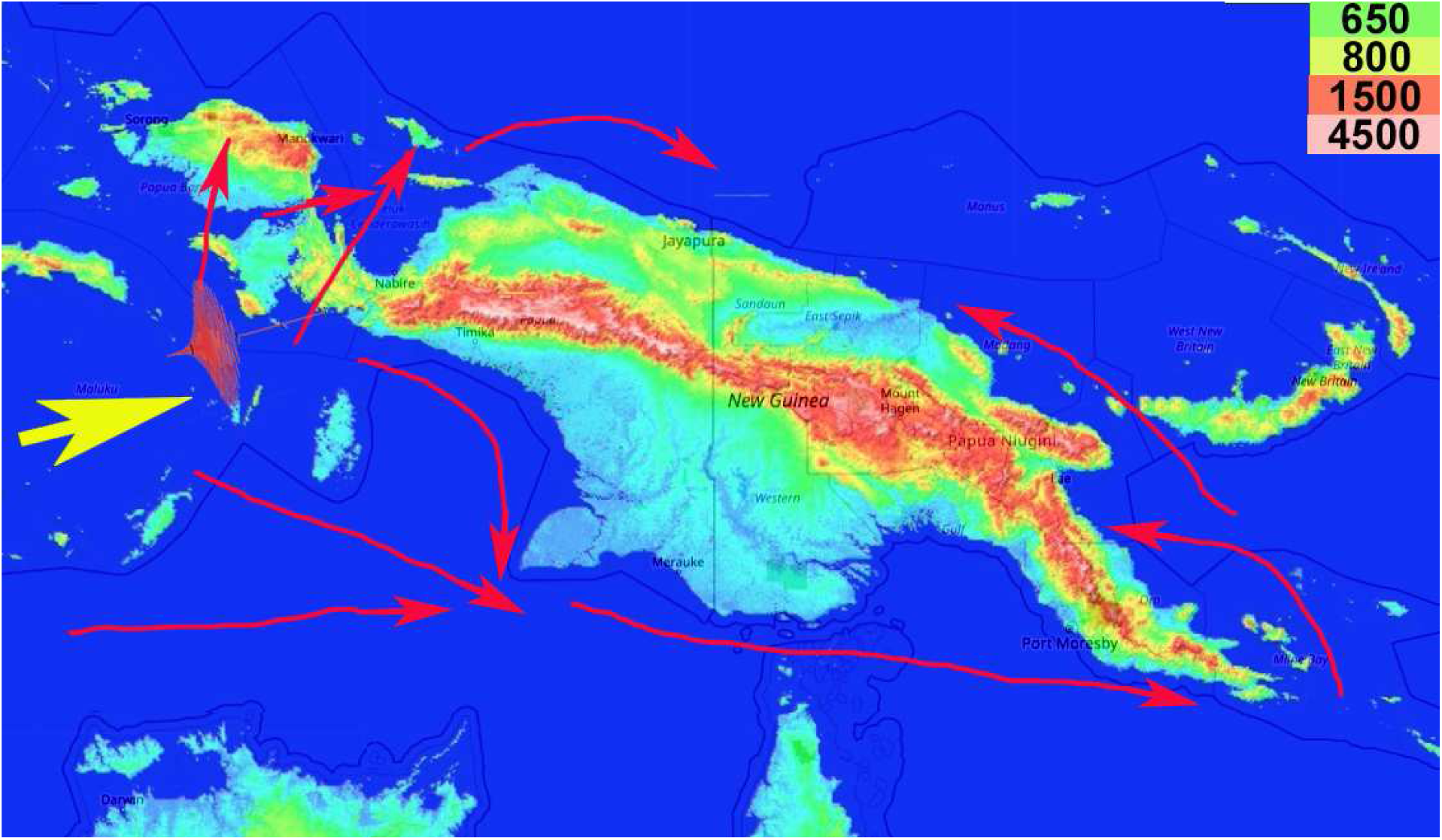
Arrival of the tsunami at New Guinea (yellow arrow) and the amplitude of the impacting waves (just beyond the main arrow). Directions of secondary and outflow waves are depicted as red arrows, showing that inundation of all but the highlands would be complete. Modern heights of mountain features are given as a heat map. Even the most conservative predictions (Table 1) indicate that all features below 800 m would be inundated.

This wave (Figure 2) would have progressed undiminished through the Java sea. Transitioning into the deeper waters of the Banda Sea (max depth 7.2km) periodicity would have decreased and this would minimize the effects of wave progression upon landfall (Figure 3). The wave would have propagated at a longer wavelength and the small islands along the northern and southern boundaries of the sea would act to focus the wave towards the Bird’s Neck region of New Guinea, reducing the wavelength and increasing wave height. The tsunami would have inundated the Bird’s Head and the southern seaboard of the island. Crossing the narrow regions of the Bird’s Neck the wave would have progressed along the northern coast of the island. The southern tip of New Guinea would have caused eddies, with a weaker (but still destructive) wave progressing up the north coast of the Island (bottom right of Figure 3).

### Effect of the Toba Supereruption on New Guinea *Saccharum* serviceability

The modelling conducted for this paper indicates that the Toba supereruption could have induced a megatsunami that would have been environmentally disastrous on the ecosystem of New Guinea. It also needs to be remembered that a tsunami was not the only consequence of the Toba supereruption.

Sugarcane needs an optimal temperature of between 20°C and 35°C (but can survive at a minimum of 15°C and a maximum of 45°C) (Ebrahim et al. 1998). Today, in New Guinea, this equates to an altitude of sea level to about 550-800m and not above 1700m (where frosts begin to happen). Though modelling of global temperatures as a result of the Toba supereruption no longer supports a ‘volcanic winter’ scenario, work by NASA and others (Robock et al. 2009) does support a significant cooling, with an average global temperature reduction of 10°C for a decade or more and a maximum temperature reduction of 20°C about five years after the reduction. Even if this only resulted in an average reduction of 4°C in the tropics (and New Guinea) this would have reduced the maximum elevation at which *Saccharum* could grow down to 250–300m (Bourke 2010).

From the modelling in this paper, the tsunami would, at a minimum, have inundated regions below 300m and below 750m at a maximum. All lowland areas below 300m, at the very least, would have been inundated and the salinity would have rendered these areas un-colonisable and un-inhabitable by plant life for decades. With the global reduction in temperature, even if *Saccharum* survived in the high valleys, they would not have survived the first cold winters. Thus a combination of a supertsunami and falling global temperatures would have eliminated *Saccharum* from New Guinea (if the genus ever existed on the island during this geological time period).

Though supporting evidence for a supertsunami impacting New Guinea 74 ka is currently sparse, Chapelle (1974) did find a transgression maximum within the reef terraces of northeast Huon, New Guinea that corresponded to this time period. If this is consistent with the Youngest Toba Event, then it supports island-wide inundation in New Guinea.

Thus, modelling indicates that the Toba supereruption may well have been a major catastrophe for the lowland regions of New Guinea.

### The Late Pleistocene Megafaunal Extinction and Sahulian Flora

During the middle to late Pleistocene (a glacial period) a major extinction event occurred that eliminated much of the megafauna of the Americas, tundral Europe and the Sahul (the single land mass formed from Australia, New Guinea and Tasmania). Current explanations for this phenomenon include: overhunting (Prideaux et al. 2009; Saltré et al. 2016); indirect effects of landscape modification (such as slash and burn farming by aboriginal peoples (Miller et al. 2005)); and the impacts of long-term climate change (Price and Webb 2006; Faith and O’Connell 2011; Dortch et al. 2016).

A full analysis of the evidence is beyond the scope of this article, particularly as most authors focus on the megafaunal extinctions themselves. However, as vegetation can be proxies for climate and human mediated change, information on the vegetative changes accompanying the late Pleistocene extinctions (the period around 45–41 ka) have started to come to light.

Work at Cuddie Springs (DeSantis et al. 2017) in Australia indicated a loss of C_4_ vegetation after ≈45 ka in southeastern Australia, coincident with increasing aridification through the middle to late Pleistocene.

The process underway (whether natural or human driven) has been described as an ‘ecosystem collapse’ (Miller et al. 2005) with egg shell data suggesting a rapid transition from a C_4_ plant diet to a C_3_ plant diet. After 45 ka, vegetation available to *Dromaius* spp (Australian emu) had become exclusively C_3_ based. There is a chicken and egg problem here… did the C_4_→C_3_ vegetative transition present one cause leading to megafaunal extinction, or was it the loss of megafaunal grazers that led to the loss of C_4_ plants? Regardless of the cause, it is clear from Australian evidence that throughout the Sahul (and this includes New Guinea) a similar process was occurring. From 58–44 ka, the abundance of plants with the C_4_ carbon fixing pathway was generally high—between 60–70%. By 43 ka, the abundance of C_4_ plants dropped to 30% and biomass burning increased. This transient shift lasted for about 3 000 years and came after the period of human colonization and directly followed megafauna extinction at 48.9–43.6 ka (Grün et al. 2010). A sudden loss of C_4_ vegetation has also been observed at the Lynch’s Crater site in northeastern Australia (Rule et al. 2012). With a more complete plant record, a dramatic drop in the occurrence of Sahulian C_4_ plants has also been reported by Lopes des Santos et al. (2013).

Megafauna extinctions occurred in New Guinea, just as they did in Australia during the Pleistocene. This would have induced a loss of C_4_ plants in New Guinea, just as happened in Australia. The resultant C_3_ scrublands would have been more fire-prone, thus reducing C_4_ containing grasslands even further. This means that New Guinea would have become increasingly inhospitable for *Saccharum* species, even if they existed on the island. The potentially dramatic increase in bushfires during this period also has implications for the interpretation of evidence interpreted as torch-burn fires due to human settlement and forest clearances.

An unforeseen consequence of the removal of magafauna (particularly large herbivores) from a region, as has been demonstrated in Africa, is the replacement of savanna by climax woodland and until the past 700 years this has been the major vegetative character of New Guinea (Malhi et al. 2016).

### Effect of the Last Glacial Maximum on New Guinea Flora

The most recent global event that significantly altered the climate was the last glacial maximum (LGM), the most proximal period at which the polar ice sheets were at their greatest extent. Growth of ice sheets commenced 33 000 years ago and maximum coverage was between 26 500 years and 19–20 000 years ago, when deglaciation commenced in the Northern Hemisphere, causing an abrupt rise in sea level. Decline of the West Antarctica ice sheet occurred between 15–14 ka, consistent with evidence for another abrupt rise in the sea level about 14 500 years ago. Ice core data, pollen data and global bog core data with isotope information have been used to create models of temperatures and vegetation patterns during this period (Clark et al. 2009; Evans et al. 2014)).

During the Last Glacial Maximum, much of the world was cold, dry, and inhospitable, with frequent storms and a dust-laden atmosphere. The dustiness of the atmosphere is a prominent feature in ice cores. Indeed, dust levels were as much as 20 to 25 times greater than now (Claquin et al. 2003). Based on the work of many authors it is possible to re-construct a sea level and vegetative biome map of Southeast Asia (Crowley and Baum 1997; Prencice et al. 2003; Wang et al. 2009). This map (Figure 4) shows the Sahul region of Australia + New Guinea + Tasmania and the Sunda region of the greater Malay peninsula. During this time Australia became increasingly arid, with New Guinea largely covered by dry forests as well as the central region of Indochina. However, India + Sri Lanka, the land bridge between Australia and New Guinea and the central Malay Peninsula, Indonesia, Borneo, Sulawesi, Sumatra, Java and the Philippines would all have been refugia for tropical C_4_ grasslands.

**Figure 4:**
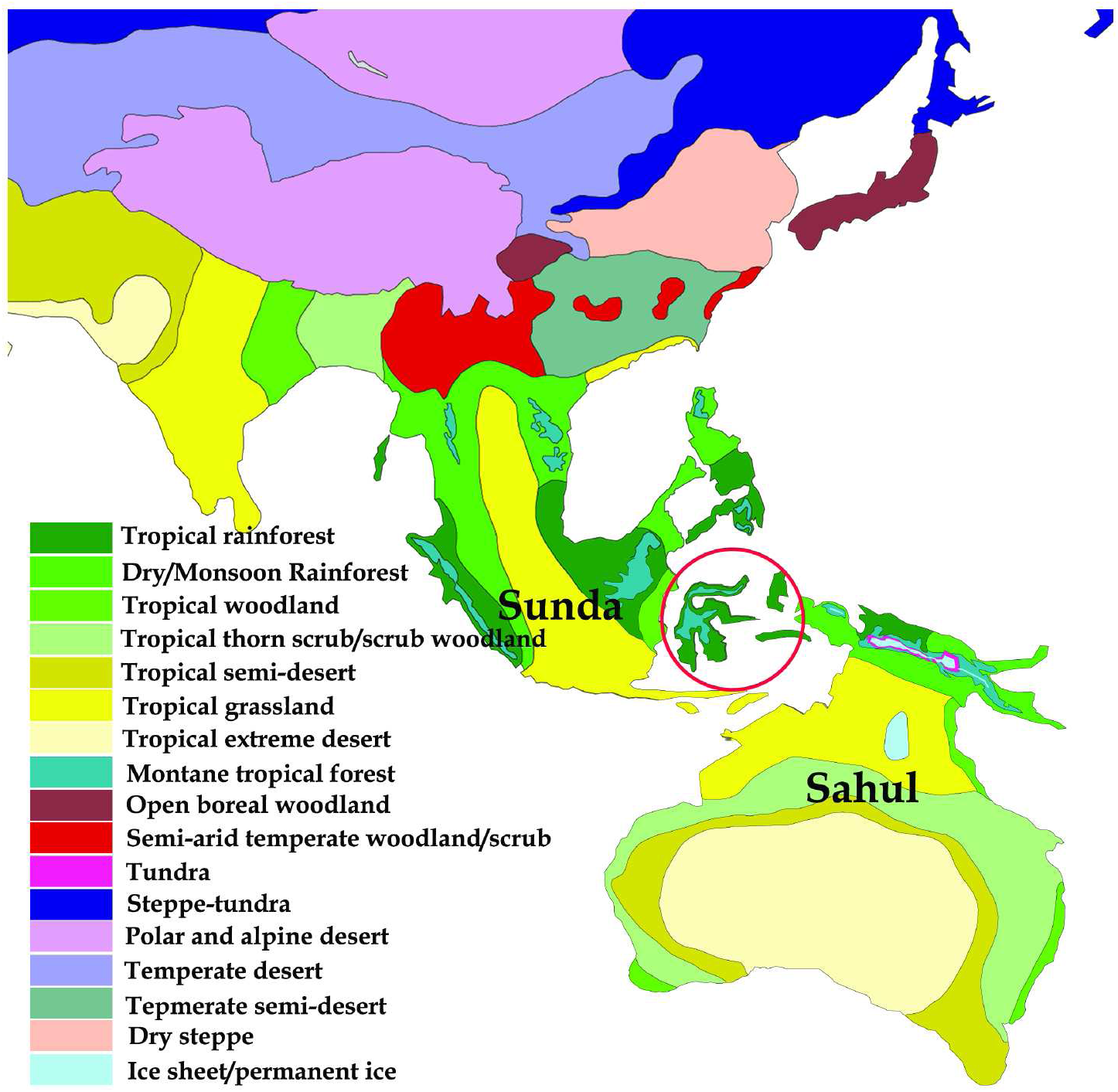
Map of south Asia and western Oceania indicating the extents of land masses during the LCM and key vegetative habitats during that period. The key gives the 17 main ecological biomes predicted for the region. The regions of the Sahul and Sunda are named. Wallacea is enclosed by a red circle.

For the first time, the specific LGM climate of New Guinea is modelled (Figure 5). The high mountains along the northern back of the island would have been covered in permanent ice with a bounding area of tundra. Next to the tundra would be montane tropical forest — cold and dry deciduous/pine tree cover up to the tree line. The majority of the island would have been dominated by dry/monsoon woodland which is too arid for *Saccharum* species to maintain a foothold. It is predicted that there was a small area of tropical rainforest at the northern coast of New Guinea. However, much of this would have been rapidly flooded as post-glacial sea levels increased. Despite adequate temperatures and high rainfall in such areas, the dense leaf cover typically means that they are unsuitable for *Saccharum* species to thrive. Thus, if the presence of *Saccharum* species in New Guinea is natural then they could only have come from Sahul… either from the Sahul itself, by island-hopping across Wallacea or by crossing from Timor into the tropical grassland of the Sahul landmass (Figure 4). However, there are no native *Saccharum* species in Australia (for this theory to be correct plants would have had to traverse from Australia via the Sahul land bridge into New Guinea). If this was the case, *Saccharum* species would have populated Australia and New Guinea, both.

**Figure 5:**
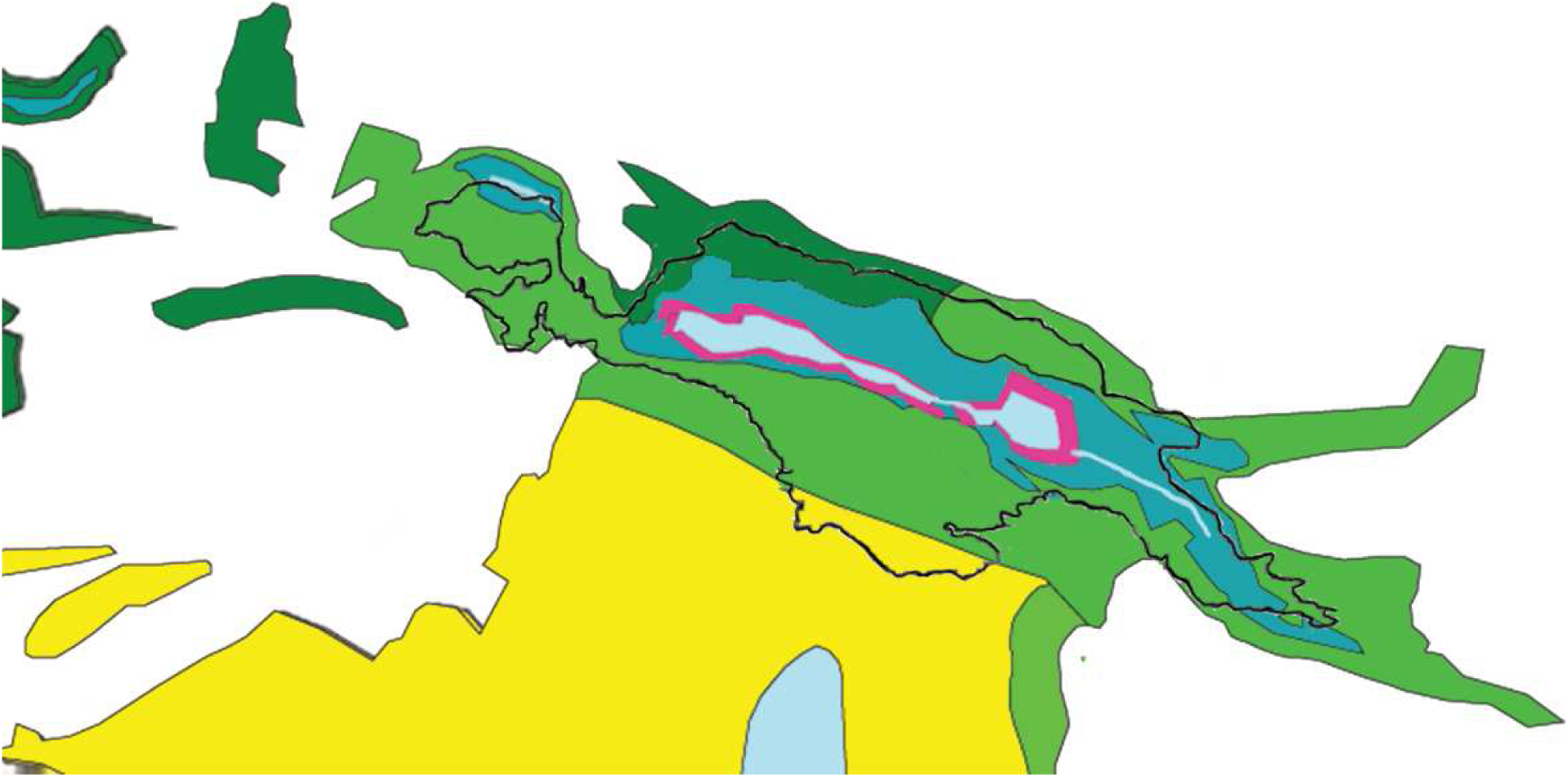
Map of New Guinea with coastline adjusted to 18 ka. The modern coastline of New Guinea is shown as a black line. To the south of New Guinea is the Sahulian land bridge and then the area of modern Australia. Most of the Sahulian land bridge is predicted to be a tropical grassland biome, with a permanent ice sheet in the centre. The model presented in this paper places permanent glaciers on the high mountains of New Guinea. These are surrounded by tundra. For a key to the colours, see Figure 4. The image clearly shows that the Sahul land mass was separated from Asia even during this time of globally lower sea levels.

If this model is wrong (and the lack of *Saccharum* species in Australia does not lend credence to the model) then *Saccharum* species would have had to enter New Guinea by another means. The islands of Wallacea (circled in Figure 4) were never connected to the remainder of the Asian or Australian continents even during the LCM and were always cut off. As the ice sheets receded and sea levels rose, they would have become even more isolated. There is also little evidence for new human migrations into New Guinea at the end of the LCM. However, as the global climate warmed and became wetter the refugia in India, south Asia and Indonesia would have allowed *Saccharum* species to expand and remain in these locales. Thus, post glaciation, it seems that *Saccharum* species that might once have been found from Mediterranean Europe through India and Indochina were now isolated in India, Indonesia, the Philippines and Borneo. The more cold tolerant *S*. *spontaneum* would have spread and diversified to the newly-warming regions.

If New Guinea had remained a home to *Saccharum* species after the LGM we would expect significant diversity of *S*. *spontaneum* there. However, this does not seem to be the case and analyses indicate that *S*. *spontaneum* is an even more recent arrival to New Guinea — yet it is distributed across the island (Grivet et al. 2004) and was previously considered to be a native species. If *S*. *spontaneum* is a recent introduction to New Guinea, then *Saccharum* as a whole cannot have evolved on the island.

## Conclusion

Since it was initially proposed by Artschwager & Brandes (1958) and up-held by Daniels and Daniels (1993) the hypothesis that sugarcane arose in New Guinea, which is a modern centre of diversity for the plant, has largely gone unchallenged. Yet, there are decades’ worth of analysis and research in the areas of Palaeoclimatology, Palaeogeography, Palaeobotany and earth sciences that have not been brought to the table. The that one of three events leading to mass extinctions at 74 ka, 45–41 ka and 16 ka makes it highly unlikely that there was an early colonization of New Guinea by *Saccharum* species. Indeed, lack of diversity of S. spontaneum makes it far more likely that *Saccharum* was a later introduction somewhere between 6–3.5 ka with the plant being introduced by Lapitan (Austronesian) peoples (Lloyd Evans and Joshi 2016).

In this respect, the cautionary tale of sweet potato (*Ipomoea batatas*) on New Guinea should be heeded. Today the sweet potato is crucial to highland New Guinea life and there are over 400 distinct cultivars on the island. Yet, no one would ever claim that the sweet potato is native to the Island. Indeed, this plant was probably domesticated in Peru about 10–8 ka (Roullier et al 2013). Whether it was brought to New Guinea by European Explorers, or by Polynesian navigators is a matter of debate, though there is no evidence of the presence of sweet potato on the island prior to 1600 CE. Indeed, an alien comparing sweet potato in New Guinea and in Peru might (wrongly) infer that the plant originated in New Guinea and was introduced to Peru. Indeed, if you ask a Papuan about the origins of sweet potato they would be adamant that it is native to the island. It is likely that the history of sugarcane on the island mirrors that of the sweet potato. It is an introduced species that, in the isolated valley microclimate and due to selective pressure from disparate tribes, has evolved and been selected to many different forms.

The ancestors of sugarcane, like many C_4_ grasses evolved in the savannas of Africa. These plants spread eastwards and evolved into Saccharum. native regions of sugarcane are India and the south China seas. However, both these regions developed commercial sugar production early and independently. As such, wild Saccharum species were replaced with commercial sugarcane cultivars prior to historical records. It is only in New Guinea, where commercial sugar production was not invented that Saccharum species only entered kitchen garden, fencing and pig fodder uses and were never commercialized on an industrial scale, that wild-looking growth of Saccharum persisted.

Our data demonstrates that Toba would have inundated areas where Saccharum grows on New Guinea 74 000 years ago killing the plant if it existed there. The last major glaciation (ending about 12 500 years ago), which happened within the period of human occupation of New Guinea would have made it impossible for sugarcane to grow on the island. Thus, modern Saccharum on New Guinea must have been introduced subsequent to this. It is likely, indeed probable, that Saccharum species were introduced along with pigs by lapitan peoples somewhere about 3-3.5 ka when they colonized coastal areas of the island. Indeed, areas where Saccharum grows remain largely congruent with those areas where Austronesian (Lapitan) languages are still spoken in New Guinea (Figure 6).

Today, pigs are a major part of the Papuan farming system; however prior to Lapitan contact there are no records of pigs on the island. Moreover, Papuan pigs posess the (Larson et al. 2007) Polynesian mitochondrial DNA marker, meaning that they originated in pensinsular southeast Asia and were introduced to New Guinea. Sugarcane is also native to this region of the Malay Pensinsula and has been used for millennia as pig fodder. It is likely, therefore, given all the arguments that *Saccharum* is not native to New Guinea that it was first introduced as pig fodder. Indeed, though some cane is used for chewing in New Guinea, members of genus *Saccharum* are associated with pigs more than any other animals. *S*. *spontaneum* is used for arrows, sugarcane hybrids are used as pig fodder and *S*. *robustum* as fencing (Lloyd Evans and Joshi 2016). Though *S*. *edule* a form of *Saccharum* with an aborted infloresence that is cooked rather like asparagus is the only form of *Saccharum* that is used solely for human consumption. However, this vegetable form of *Saccharum* is consumed throughout Wallacea and Malaysia and may have been introduced during the period of Malay contact with New Guinea (c. 1500–1900 CE).

Just like pigs, which have become feral on the island, *Saccharim* are introduced species in New Guinea, which have become feral. Despite agriculture being truly ancient in New Guinea, farming that could support sugarcane growth did not appear until about 4.5 ka. Moreover, egetation patterns not dissimilar to those of today seem to have become established rather widely by 4 ka. This involved the expansion of grassland at the expense of secondary forest (Haberle 1994). In the context of Kuk, Golson and Gardner (1990) discuss the evidence for the appearance of three features of modern highlands agricultural technology thought to have been developed in response to the late Holocene environmental transformation from forest to grassland: soil tillage from 2 ka, tree fallowing from ca. 3.2 ka and raised-bed cultivation by ca. 0.4 ka This would make farming systems that support *Saccharum* species rather late developments in the New Guinea highlands as they extend from Lapitan settlements in the lowlands.

The data presented in this paper make it highly unlikely that *Saccharum* as a genus evolved on New Guinea. Though *Miscanthus* is present, the more immediate sister species *Miscanthidium* and *Narenga* are not. Moreover, three independent cataclysms would have eliminated *Saccharum* from the island. Whilst it is possible that S. spontaneum might have survived in the dry grasslands of the Last Glacial Maximum, *S*. *officinarum* could not and the low genetic diversity in *S*. *spontaneum* speaks to it being a more recently introduced species. The currently accepted model of the origins of sugarcane needs to be re-visited in light of these findings.

## Materials and Methods

### Tsunami Modelling

The inception phase for a supervolcano derived megatsunami is the fall-back of ejecta from the initial caldera explosion. In many ways, this echoes the tsunami inception from a meteor impact, which is well studied. We start from the perfect conditions. In an uniform ocean of depth *h*, an initial displacement 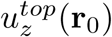 and velocity 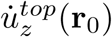 at the sea surface evolves into vector tsunami waveforms at point r of 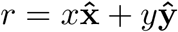, depth *z* and time *t* (from Ward (2002)).

A fully three-dimensional waveform can be written as:

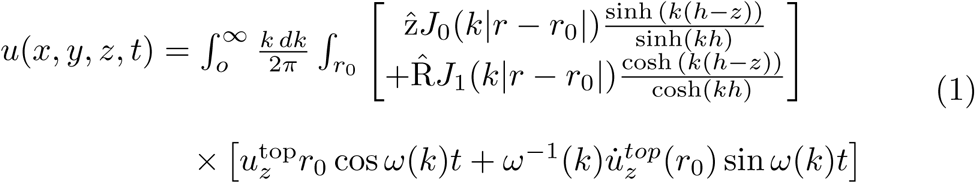

where 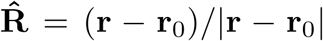, wavenumber *k* = |*k*|, the frequency 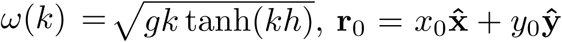, *dk* = *dk*_*x*_*dk*_*y*_, 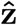 is the unit vector **z***/*|**z**|, *d***r**_0_ = *dx*_0_*dy*_0_; *g* is the gravitational constant (9.8 ms^-1^) and the *J*_*n*_ represent Bessel functions. Equation 1 is applicable to an oceanic disturbance due to any root cause and appropriate initial conditions need to be chosen. Impact tsunami can, however, be approximated by cavitation effects (the effect seen from dropping any object into water). At the final stage of a cavitation event motion transitions from a mostly downward to a mostly upward vector, leading to the initiation and propagation of the tsunami wave. Ward & Asphaug (2000) assumed that, at this stage, the vertical surface velocity to be negligible (ie *x* → 0) and the surface displacement follows a parabola:

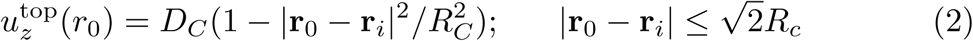

where *D*_*C*_ and *R*_*C*_ are, respectively, the depth and radius of the cavity. 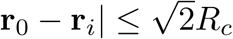 expresses the assumption that the volume of the cavity equals the volume of water deposited on the rim. Entering these assumptions into (1) and setting *z* = 0 we obtain:

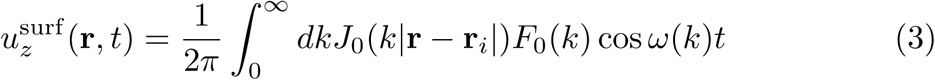

where

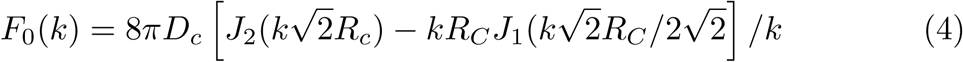

real oceans are non-uniform in depth and the Toba tsunami would have traversed several deep/shallow ocean bed depth changes. Equation (3) can be generalized to non uniform depth oceans by calculation of the raypath-specific travel time *T* (*ω*, **r**, **r**_*i*_) and incorporating the shoaling factors *G*(**r**, **r**_*i*_) and *S*_*L*_(*ω*, **r**, **r**_*i*_) (see Ward (2001) for further details. Adding these terms to (3) and changing the wavenumber to frequency (this is constant as waves traverse water of different depth) we get:

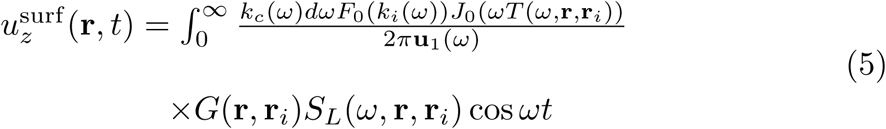

For most (but not all) continental shelf encounters shoaling tsunami waves will grow according to Equation 5 until a point r_*c*_ is reached where the wave height *A*(**r**_*c*_) equals a given fraction of the ocean depth *ψh*(**r**_*c*_). Typically this critical height is equal to the eventual run-up height *R* at the shore. The run-up correction is thus given as:

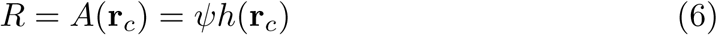

if *A*(**r**) is the amplitude of a pack of waves in water of depth *h*(**r**) then, entering these values in the shoaling factor:

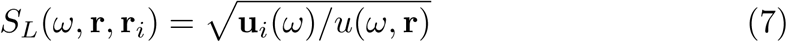

then the wave amplitude in shallow water becomes:

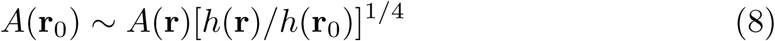

thus it follows that the depth *h*(**r**_*c*_) where *A*(**r**_0_) = *ψh*(**r**_0_) is *h*(**r**_*c*_) = *ψ*^1*/*5^*A*(**r**)^4*/*5^*h*(**r**)^1*/*5^; thus:

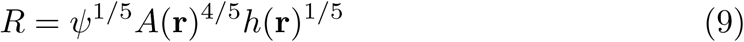

Returning to equation (2), this assumes that the wave propagates in all directions. However, in a channel this is not possible. Wave propagation onto shore will be significantly less that the wave propagation at the open ends of the channel. Thus, in the case of the Toba tsunami the assumption that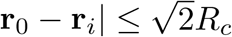 is not valid. As the water was dissipated through a channel, one where the easterly end is larger than the western end means that the wave height can be greater than the depth of water. Thus a scale factor is applied:

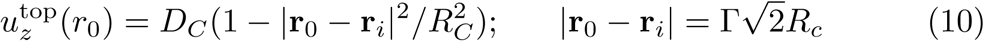

Γ is varied in intervals of 0.2 from 1 to 1.6.

### Model Implementation

Models were implemented in Wolfram Mathematica (Wolfram Research) version 10.2.2 run on a Ubuntu 18.4 running on a 32GiB memory, 2TiB hard drive laptop. After initial testing and implementation simulations were run with varying sea floor depths and at varying Γ correction factors. Wave propagation in the straits of Molucca were also modelled as a time series.

### Vegetation Mapping for the Last Global Maximum in East Asia and Australasia

Methodology essentially followed that of Ray and Adams (2001) with coastal regions assumed to be 120m extended from present day coastal limits. As this study focusses on East Asia and Australasia, the effect of ice sheets is generally minimized. The exception being New Guinea, where the high mountains would have generated their own climate. The information on glaciation from Peterson et al. (2004) was used to generate a more detailed biome map of New Guinea.

Maps and features were drawn with the Linux package QGIS 3.12, with annotation finished in PhotoShop. Coastal maps were those of Ray and Adams (2001) imported into QGIS and corrected to map coordinates based on QGIS internal maps. Separate maps of Eurasia and Australasia were combined into a single map. A separate map for New Guinea and the Sahulian land bridge to Australia was produced. The data of Ray and Adams (2001) were taken for Indochina and South East Asia, but the data for Sahul (particularly New Guinea) were updated based on archaeological and bog strata evidence (Van der Kaars 1991; Tudhope et al. 1995; Miller et al. 2012; Beehler & Laman 2020).

